# Inositol 1,4,5-trisphosphate receptor type2-independent Ca^2+^ release from the endoplasmic reticulum in astrocytes

**DOI:** 10.1101/321042

**Authors:** Yohei Okubo, Kazunori Kanemaru, Junji Suzuki, Kenta Kobayashi, Kenzo Hirose, Masamitsu Iino

**Affiliations:** Department of Pharmacology, Graduate School of Medicine, The University of Tokyo, Tokyo 133-0033, Japan; Department of Cellular and Molecular Pharmacology, Nihon University School of Medicine, Tokyo 173-8610, Japan.; Department of Physiology, University of California San Francisco, San Francisco, CA 94158, USA.; Section of Viral Vector Development, National Institute for Physiological Sciences, Okazaki 444-8585, Japan.; The Graduate University for Advanced Studies (SOKENDAI), Hayama 240-0193, Japan.; Department of Neurobiology, Graduate School of Medicine, The University of Tokyo, Tokyo 133-0033, Japan

**Keywords:** Astrocyte, Endoplasmic reticulum, Mitochondria, Calcium, Inositol 1,4,5-trisphosphate

## Abstract

Accumulating evidence indicates that astrocytes are actively involved in the physiological and pathophysiological functions of the brain. Intracellular Ca^2+^ signaling, especially Ca^2+^ release from the endoplasmic reticulum (ER), is considered to be crucial for the regulation of astrocytic functions. Mice with genetic deletion of inositol 1,4,5-trisphosphate receptor type 2 (IP_3_R2) are reportedly devoid of astrocytic Ca^2+^ signaling, and thus widely used to explore the roles of Ca^2+^ signaling in astrocytic functions. While functional deficits in IP_3_R2-knockout (KO) mice have been found in some reports, no functional deficit was observed in others. Thus, there remains a controversy regarding the functional significance of astrocytic Ca^2+^ signaling. To address this controversy, we re-evaluated the assumption that Ca^2+^ release from the ER is abolished in IP_3_R2-KO astrocytes using a highly sensitive imaging technique. We expressed the ER luminal Ca^2+^ indicator G-CEPIA1*er* in cortical and hippocampal astrocytes to directly visualize spontaneous and stimulus-induced Ca^2+^ release from the ER. We found attenuated but significant Ca^2+^ release in response to application of norepinephrine to IP_3_R2-KO astrocytes. This IP_3_R2-independent Ca^2+^ release induced only minimal cytosolic Ca^2+^ transients but induced robust Ca^2+^ increases in mitochondria that are frequently in close contact with the ER. These results indicate that ER Ca^2+^ release is retained and is sufficient to increase the Ca^2+^ concentration in close proximity to the ER in IP_3_R2-KO astrocytes.

## Introduction

Accumulating evidence indicates that astrocytes play not only trophic and supportive roles but also active roles in the physiological and pathophysiological functions of the brain (Volterra & Meldolesi, 2005; Barres, 2008). Therefore, clarifying the mechanisms for regulation of astrocytic functions has become a crucial issue in neuroscience. Intracellular Ca^2+^ signaling has attracted attention because astrocytes show robust spontaneous and stimulus-induced Ca^2+^ transients (Verkhratsky, Orkand, & Kettenmann, 1998; Bazargani & Attwell, 2016).

Ca^2+^ release from the endoplasmic reticulum (ER) via inositol 1,4,5-trisphosphate receptor (IP_3_R) has been recognized as a major component of spontaneous and G_q_-coupled receptor-induced Ca^2+^ signaling in astrocytes. Therefore, to explore the significance of astrocytic Ca^2+^ signaling, mice with genetic deletion of IP_3_R type 2 (IP_3_R2) (Li et al., 2005), which is enriched in astrocytes (Sharp et al., 1999; Holtzclaw et al., 2002; Zhang et al., 2014), have been widely used. In IP_3_R2-knockout (KO) mice, Ca^2+^ transients in astrocytes were considered to be absent based on an imaging study of cytosolic Ca^2+^ transients by fluorescent Ca^2+^ indicator dyes (Petravicz, Fiacco, & McCarthy, 2008). Physiological and pathophysiological phenotypes have been observed in IP_3_R2-KO mice, including plasticity and learning (Takata et al., 2011; Chen et al., 2012; Navarrete et al., 2012; Perez-Alvarez et al., 2014; Padmashri, Suresh, Boska, & Dunaevsky, 2015; Kim et al., 2016; Monai et al., 2016; Yang et al., 2016), homeostasis of K^+^ (Wang et al., 2012a; b), pathogenesis of stroke, traumatic brain injury and neurodegenerative disease (Dong, He, & Chai, 2013; Kanemaru et al., 2013; Li et al., 2015; Rakers & Petzold, 2016; Saito et al., 2018), and depression-like behaviors (Cao et al., 2013). However, several studies have reported no changes in basal synaptic activity and behavior (Petravicz, Fiacco, & McCarthy, 2008; Agulhon et al., 2013; Petravicz, Boyt, & McCarthy, 2014), hippocampal synaptic plasticity (Agulhon, Fiacco, & McCarthy, 2010), and neurovascular coupling (Nizar et al., 2013; Takata et al., 2013; Bonder & McCarthy, 2014) in IP_3_R2-KO mice. These contradictory results have caused confusion about the functional significance of Ca^2+^ signaling in astrocytes (Hamilton & Attwell, 2010; Fiacco & McCarthy, 2018; Savtchouk & Volterra, 2018).

One of the possible explanations that can resolve this controversy is that IP_3_R2-independent Ca^2+^ signaling exists and contributes to the functions of IP_3_R2-KO astrocytes. In fact, recent studies using genetically encoded Ca^2+^ indicators (GECIs) have clearly shown the presence of Ca^2+^ signals in IP_3_R2-KO astrocytes (Kanemaru et al., 2014; Srinivasan et al., 2015; Rungta et al., 2016; Agarwal et al., 2017). These IP_3_R2-independent Ca^2+^ transients were thought to be mediated by Ca^2+^ influx via the plasma membrane (Shigetomi et al., 2012; Srinivasan et al., 2015; Rungta et al., 2016) or Ca^2+^ release from mitochondria (Agarwal et al., 2017). These studies suggest that increases in the concentration of Ca^2+^ originating from sources other than the ER may regulate the functions of astrocytes. However, it remains elusive whether ER Ca^2+^ release is completely abolished in IP_3_R2-KO astrocytes.

We thus examined whether IP_3_R2-independent Ca^2+^ release from the ER is present in IP_3_R2-KO astrocytes. Because cytosolic Ca^2+^ transients can be generated by Ca^2+^ derived from sources other than the ER, we directly detected ER Ca^2+^ release as a decrease in the Ca^2+^ concentration within the ER ([Ca^2+^]_ER_). We recently developed a series of calcium-measuring organelle-entrapped protein indicators (CEPIAs) that allow visualization of Ca^2+^ dynamics within the ER and mitochondria with a high signal-to-noise ratio (Suzuki et al., 2014; Okubo et al., 2015; Suzuki, Kanemaru, & Iino, 2016). In this study, we expressed the ER Ca^2+^ indicator G-CEPIA1*er* in cortical and hippocampal astrocytes of IP_3_R2-KO mice. We found attenuated but significant Ca^2+^ release from the ER. We also used the mitochondrial Ca^2+^ indicator CEPIA2*mt* and successfully observed robust Ca^2+^ transfer into mitochondria from the ER in IP_3_R2-KO astrocytes. These findings allowed us to reinterpret the results from IP_3_R2-KO mice and shed new light on the significance of astrocytic Ca^2+^ signaling.

## Materials and Methods

### Animals

All animal experiments were carried out in accordance with the regulations and guidelines of the Institutional Animal Care and Use Committee at The University of Tokyo and were approved by the Institutional Review Committee of the Graduate School of Medicine, The University of Tokyo. C57BL/6 mice were used as wild-type (WT) mice. IP_3_R2-KO mice (Li et al., 2005) were obtained from J. Chen (University of California at San Diego). Mice were kept under a 12 h light/dark cycle with *ad libitum* access to food and drinking water.

### Preparation of viral vectors

To generate adeno-associated viruses (AAVs) for astrocyte-specific expression of GECIs, the cytomegalovirus promoter of pAAV-MCS (AAV Helper Free Expression System, Cell Biolabs, Inc., San Diego, CA) was replaced with the gfaABC_1_D astrocyte-specific promoter derived from pZac2.1-gfaABC1D-Lck-GCaMP3 (Shigetomi et al., 2013). G-CEPIA1*er*, CEPIA2*mt*, GCaMP6f, and ER-YFP were inserted downstream of the gfaABC_1_D promoter to generate pAAV-gfaABC_1_D-G-CEPIA1*er*, pAAV-gfaABC_1_D-CEPIA2*mt*, pAAV-gfaABC_1_D-GCaMP6f, and pAAV-gfaABC_1_D-ER-YFP. AAV5 vectors were packaged using the AAV Helper Free Expression System. The packaging plasmids (pAAV-RC5 and pHelper) and transfer plasmid (pAAV-gfaABC_1_D-G-CEPIA1*er*, pAAV-gfaABC_1_D-CEPIA2*mt*, pAAV-gfaABC_1_D-GCaMP6f, or pAAV-gfaABC_1_D-ER-YFP) were transfected into HEK293T cells using the calcium phosphate method. The medium was replaced at 18 h after transfection with fresh medium, and the cells were incubated for 48 h. Harvested cells were lysed by repeated freezing and thawing, and a crude cell extract containing AAV5 vector particles was obtained. AAV5 vector particles were purified by ultracentrifugation with cesium chloride. The purified particles were dialyzed against PBS and then concentrated by ultrafiltration using an Amicon 10K MWCO filter (Merck Millipore, Darmstadt, Germany). The copy number of the viral genome (vg) was determined by real-time quantitative PCR.

### AAV injection

Male C57BL/6 mice or IP_3_R2-KO mice (postnatal day 56–120) were anesthetized with isoflurane (induction at 5%, maintenance at 1–2%, vol/vol, MK-A100, Muromachi, Kyoto, Japan). The mice were placed in a stereotaxic frame (SR-5M-HT, Narishige, Tokyo, Japan). The skull was thinned (about 1 mm in diameter) above the right parietal cortex using a burr powered by a high speed drill (ULTIMATE XL-D, NSK, Kanuma, Japan).

AAV5-gfaABC_1_D-G-CEPIA1*er* (0.98 or 1.3 × 10^13^ vg ml^−1^), AAV5-gfaABC_1_D-CEPIA2*mt* (0.98 × 10^13^ vg ml^−1^), AAV5-gfaABC_1_D-GCaMP6f (1.1 × 10^13^ vg ml^−1^), or pAAV-gfaABC_1_D-ER-YFP (3.0 × 10^13^ vg ml^−1^) was unilaterally injected into the cortex (1.5–2 mm posterior to the bregma, 1–1.5 mm lateral to the midline, and 300 μm from the surface) or hippocampus (2 mm posterior to the bregma, 1.5 mm lateral to the midline, and 1.6 mm from the surface) through glass pipettes. A viral solution (1 μL) was delivered at a rate of 200 nL min^−1^ using a micropump (Legato 130, KD scientific, Holliston, MA). Glass pipettes were left in place for at least 10 min. Mice were sacrificed at 14–28 days after AAV injection for imaging and immunohistochemistry.

### Preparation of brain slices and imaging

Coronal cortical or hippocampal slices (300 μm in thickness) were prepared as described previously (Edwards, Konnerth, Sakmann, & Takahashi, 1989). Slices were prepared in ice-cold artificial cerebrospinal fluid (ACSF) bubbled with 95% O_2_ and 5% CO_2_ using a vibrating slicer (PRO7, Dosaka, Kyoto, Japan). Slices were incubated in a holding chamber containing ACSF bubbled with 95% O_2_ and 5% CO_2_ at 35°C for 1 h and then returned to 23°C. ACSF contained (in mM) 125 NaCl, 2.5 KCl, 2 CaCl_2_, 1 MgSO_4_, 1.25 NaH_2_PO_4_, 26 NaHCO_3_, and 20 glucose. Imaging was carried out with a two-photon microscope (TSC MP5, Leica, Wetzlar, Germany) equipped with a water immersion objective (×25, NA 0.95, HCS IR APO, Leica) and Ti:sapphire laser (MaiTai DeepSee; Spectra Physics, Santa Clara, CA). Slices were transferred to a recording chamber under a microscope and continuously perfused with ACSF bubbled with 95% O_2_ and 5% CO_2_. Tetrodotoxin (1 μM) was added to ACSF to inhibit neuronal activities throughout imaging. Norepinephrine (NE, 10 μM) and cyclopiazonic acid (CPA, 50 μM) were dissolved in ACSF and administered through the perfusion system of the recording chamber. The excitation wavelength was 900–920 nm. Emitted fluorescence was filtered by a 500–550 nm barrier filter and detected with photomultiplier tubes. Data were acquired in time lapse XY-scan mode (0.2–0.5 Hz). Experiments were carried out at room temperature (22–24°C).

### Data analysis

Data were analyzed using ImageJ software. Astrocytes were analyzed in layer 2/3 of the cortex or in the CA1 stratum radiatum of the hippocampus. Fluorescence intensities were corrected for background fluorescence by measuring a non-fluorescent area and normalized by the average of the first 10 or 20 frames to calculate the fractional changes in fluorescence intensity (Δ*F*/*F*_0_). Spontaneous G-CEPIA1*er* responses were manually detected and analyzed. Fluorescence changes with a rate of decrease faster than −0.015 Δ*F*/*F*_0_ s^−1^ and amplitude larger than −0.05 Δ*F*/*F*_0_ were defined as responses. Synchronous decreases in G-CEPIA1*er* fluorescence throughout a single astrocyte were defined as global responses. Localized fluorescence decreases with about 10–15 μm in diameter at the process region were defined as process-localized responses. The amplitude of spontaneous G-CEPIA1*er* responses was measured after smoothing time courses of Δ*F*/*F*_0_ by a moving average with five frame windows. The amplitude of NE-induced G-CEPIA1*er*, CEPIA2*mt*, GCaMP6f, or ER-YFP responses was defined as the average of Δ*F*/*F*_0_ within the 3-min time window starting from the start of NE application. The amplitude of the CPA-induced G-CEPIA1*er* fluorescence decrease was defined as the Δ*F*/*F*_0_ value in a steady state. For statistical analyses, Student’s *t*-test was performed. The asterisks in the figures indicate the following: \**P* < 0.05, \*\**P* < 0.01, and \*\*\**P* < 0.001.

## Results

### G-CEPIA1*er* detects Ca^2+^ release from the ER in cortical astrocytes

We used a serotype 5 AAV carrying the minimal astrocyte-specific gfaABC_1_D promoter to express G-CEPIA1*er* in astrocytes of the adult mouse cortex (Shigetomi et al., 2013). G-CEPIA1*er*-expressing astrocytes in acute cortical slice preparations were imaged using a two-photon microscope (Fig. 1A). At subcellular levels, expression of G-CEPIA1*er* was observed throughout the soma and processes (Fig. 1A). This G-CEPIA1*er* distribution was consistent with the distribution pattern of the ER in astrocytes (Patrushev, Gavrilov, Turlapov, & Semyanov, 2013).

**Figure 1.**
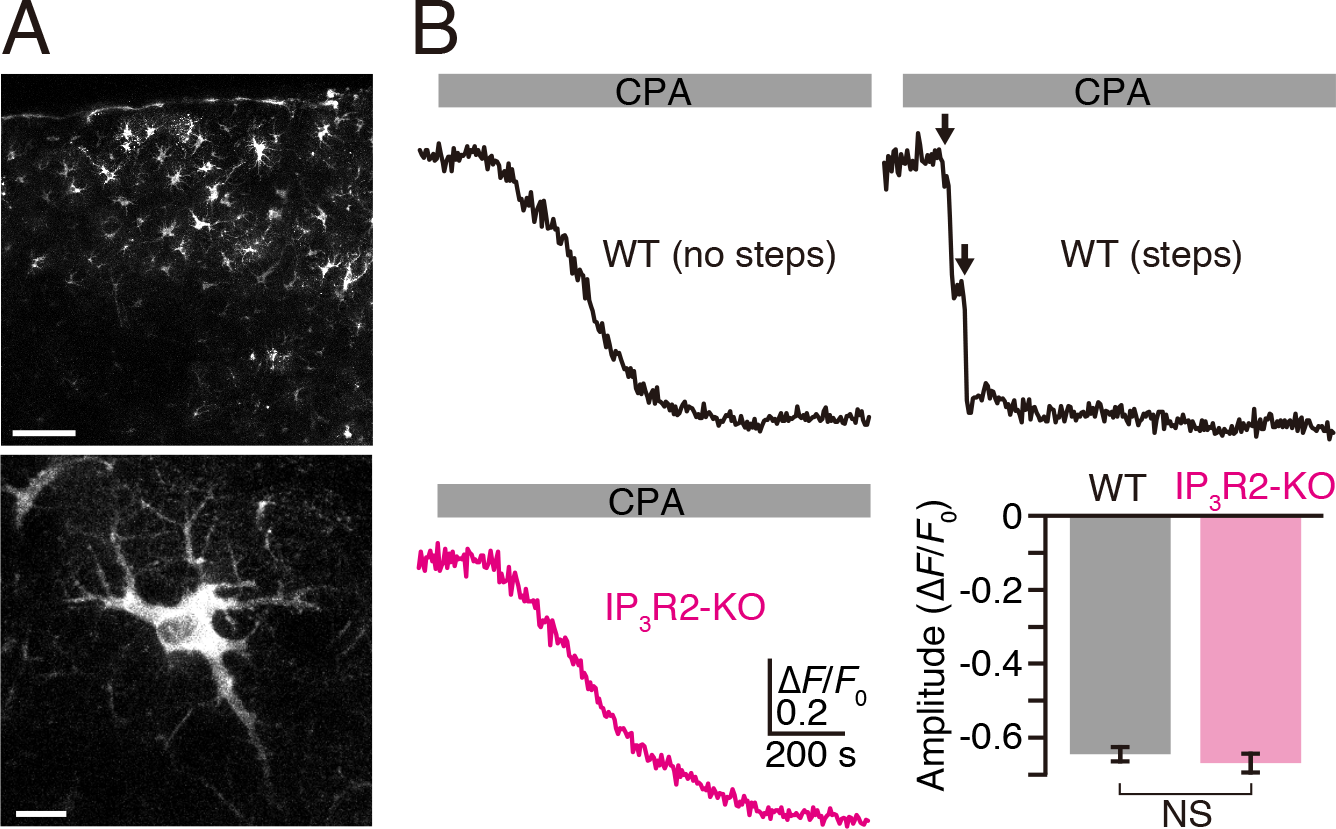
Ca^2+^ imaging within the ER of cortical astrocytes using G-CEPIA1*er*. (A) G-CEPIA1*er*-expressing astrocytes in the acute cortical slices. Maximum intensity projection of 22 (top) and 7 (bottom) images acquired by 1-μm interval. Scale bars, 100 μm (top) and 10 μm (bottom). (B) SERCA inhibitor-induced Ca^2+^ depletion confirmed that G-CEPIA1*er* reflected [Ca^2+^]_ER_. Representative Δ*F*/*F*_0_ traces of G-CEPIA1*er* upon application of CPA (50 μM, gray bar) to WT (black traces) and IP_3_R2-KO (magenta trace) astrocytes. Majority of WT astrocytes (17 cells among 23 cells) showed a stepwise decrease in G-CEPIA1*er* fluorescence as indicated by arrows in the top right trace. The rest of WT astrocytes (6 cells among 23 cells, top left) and all IP_3_R2-KO astrocytes (21 cells, bottom left) showed a gradual decrease in G-CEPIA1*er* fluorescence. The region of interest to measure Δ*F*/*F*_0_ covered the entire cell. Bottom right: summary of the amplitude of G-CEPIA1*er* fluorescence decrease upon application of CPA (*n* = 23 cells for WT and 21 cells for IP_3_R2-KO, mean ± s.e.m.). No significant difference was detected between WT and IP_3_R2-KO (*P* = 0.447, unpaired Student’s *t*-test).

G-CEPIA1*er*-expressing layer 2/3 astrocytes were imaged to analyze Ca^2+^ release from the ER. We first applied CPA, a sarco/endoplasmic reticulum Ca^2+^-ATPase (SERCA) inhibitor, to confirm G-CEPIA1*er* responses to [Ca^2+^]_ER_ decreases. Inhibition of SERCA-dependent Ca^2+^ uptake results in Ca^2+^ depletion of the ER. Indeed, CPA treatment resulted in a marked decrease in the G-CEPIA1*er* fluorescence intensity of both WT and IP_3_R2-KO astrocytes, confirming that G-CEPIA1*er* reflects [Ca^2+^]_ER_ (Fig. 1B). There was no significant difference in the amplitudes of the CPA-induced G-CEPIA1*er* fluorescence decrease between WT and IP_3_R2-KO astrocytes, indicating that the basal [Ca^2+^]_ER_ in IP_3_R2-KO astrocytes was similar to that in WT astrocytes (Fig. 1B). However, the rate of decrease in [Ca^2+^]_ER_ was often greater in WT astrocytes than in IP_3_R2-KO astrocytes (Fig. 1B). In the majority of WT astrocytes (17 cells among 23 cells), [Ca^2+^]_ER_ decreased in a stepwise manner as shown in Fig. 1B. Each step appeared to reflect spontaneous Ca^2+^ release events that are described in the following (Fig. 3). In the remaining WT astrocytes (6 cells among 23 cells) and all of IP_3_R2-KO astrocytes (21 cells) showed a gradual decrease in [Ca^2+^]_ER_ due to Ca^2+^ leakage from the CPA-treated ER (Carreras-Sureda, Pihán, & Hetz, 2018).

### Ca^2+^ release from the ER is retained in IP_3_R2-KO astrocytes

We next investigated G_q_-coupled receptor-induced Ca^2+^ release from the ER in astrocytes. Norepinephrine (NE), which is released from the axons of locus coeruleus neurons, induces cell-wide Ca^2+^ release from the ER through activation of α_1_-adrenergic receptors and following IP_3_ signaling in WT astrocytes *in vivo* (Ding et al., 2013; Paukert et al., 2014; Srinivasan et al., 2015). Bath application of NE (10 μM) induced a large decrease in [Ca^2+^]_ER_ throughout the soma and processes of WT astrocytes, indicating robust and cell-wide Ca^2+^ release from the ER (Fig. 2A and C). Following washout of NE, there was slow recovery of [Ca^2+^]_ER_ due to Ca^2+^ reuptake by the ER. The time course was consistent with ER Ca^2+^ dynamics induced by IP_3_R-mediated Ca^2+^ release in other cell types shown in our previous studies (Suzuki et al., 2014; Okubo et al., 2015). In some WT astrocytes, [Ca^2+^]_ER_ undershot the dynamic range of G-CEPIA1*er* (Fig. 2A and C). In such cases, response amplitude would be underestimated. Most GECIs including G-CEPIA1*er* are sensitive to pH. To exclude the possibility that the observed changes in the G-CEPIA1*er* fluorescence intensity was influenced by a change in pH within the ER, we expressed ER-localized YFP that is sensitive to pH but insensitive to Ca^2+^ (Suzuki et al., 2014). ER-YFP showed no response to NE, confirming that NE-induced G-CEPIA1*er* responses reflected decreases in [Ca^2+^]_ER_ (Fig. 2C).

**Figure 2.**
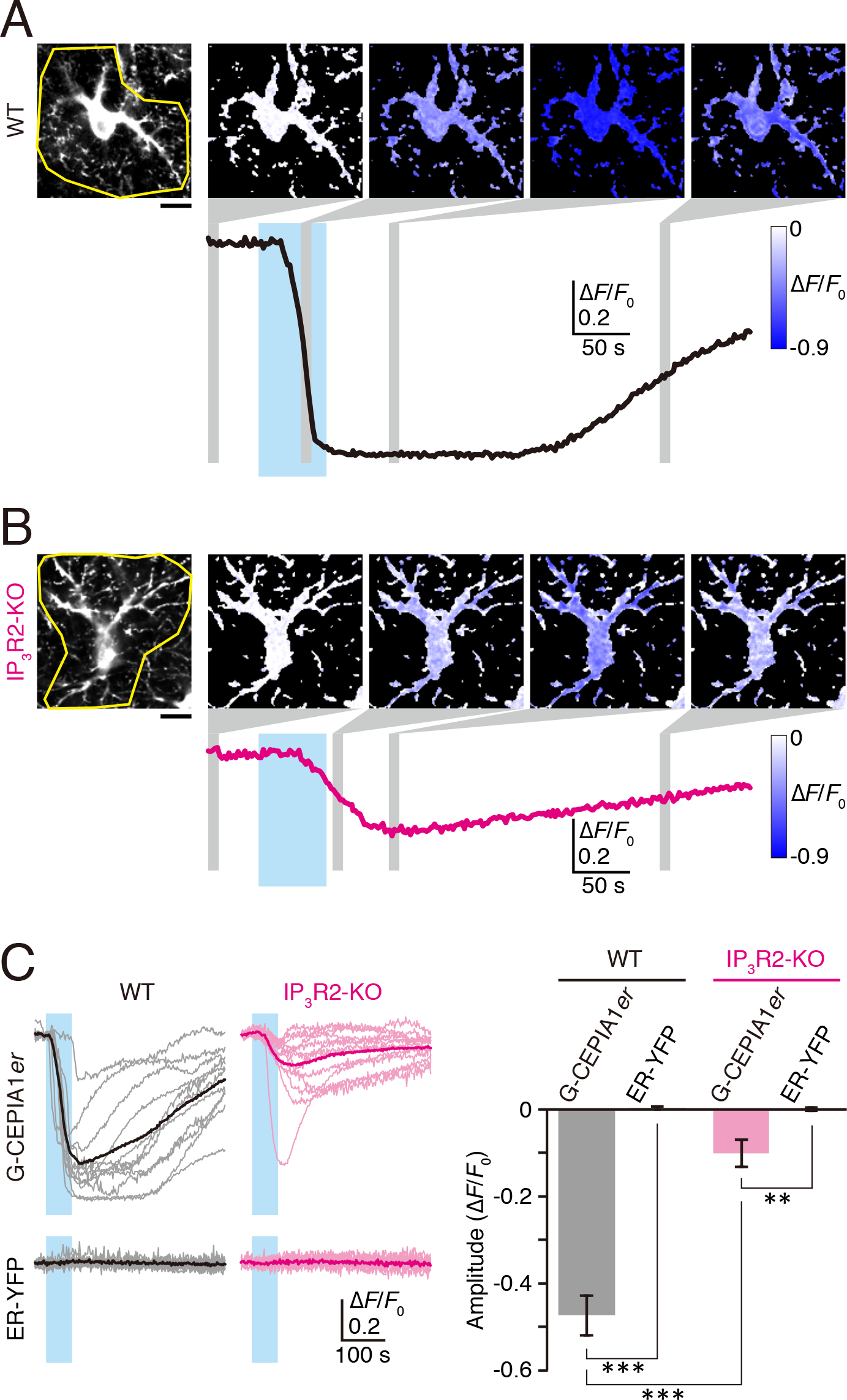
IP_3_R2-independent Ca^2+^ release in cortical astrocytes. (A and B) Representative G-CEPIA1*er* responses upon application of NE (10 μM, cyan bars) to WT astrocytes (A) and IP_3_R2-KO astrocytes (B) in cortical slices. The yellow circles in the leftmost images indicate the region of interest for the Δ*F*/*F*_0_ traces. Pseudocolor images indicate the average of five consecutive Δ*F*/*F*_0_ images within the time zones indicated by gray vertical bars. Scale bars, 10 μm. (C) Left: G-CEPIA1*er* and ER-YFP responses upon application of NE (10 μM, cyan bars) to WT (black) and IP_3_R2-KO (magenta) astrocytes. Individual Δ*F*/*F*_0_ traces (pale color) and average traces (deep color) are shown. Right: summary of the response amplitudes of Δ*F*/*F*_0_ traces (mean ± s.e.m.). For G-CEPIA1*er*, *n* = 11 cells for WT and 13 cells for IP_3_R2-KO. For ER-YFP, *n* = 12 cells for WT and 10 cells for IP_3_R2-KO. Asterisks indicate the statistical significance. \*\**P* < 0.01, \*\*\**P* < 0.001, unpaired Student’s *t*-test.

We then studied the ER Ca^2+^ dynamics in IP_3_R2-KO astrocytes. Although previous studies have shown that NE-induced Ca^2+^ release is absent in IP_3_R2-KO astrocytes, NE application induced a smaller but significant decrease in [Ca^2+^]_ER_ throughout the soma and processes of IP_3_R2-KO astrocytes, while there was no change in ER-YFP fluorescence (Fig. 2B and C). Thus, contrary to the previous assumption, this observation clearly indicates that ER Ca^2+^ release is retained in IP_3_R2-KO astrocytes.

During our observation of [Ca^2+^]_ER_ in WT astrocytes, we found spontaneous decreases in [Ca^2+^]_ER_ (Fig. 3). There were two types of spontaneous events: a “global” response that showed a synchronous decrease in [Ca^2+^]_ER_ throughout the astrocyte (Fig. 3A), and a “process-localized” response that was localized at processes with diameters of about 10–15 μm (Fig. 3B). However, we did not find either “global” or “process-localized” responses in IP_3_R2-KO astrocytes (Fig. 3C). Thus, in IP_3_R2-KO astrocytes, spontaneous Ca^2+^ release events, if any, appeared to be attenuated to a level undetectable by G-CEPIA1*er*. These spontaneous Ca^2+^ release dynamics may at least partly correspond to cell-wide and localized Ca^2+^ signals imaged by cytosolic Ca^2+^ indicators in previous studies (Kanemaru et al., 2014; Srinivasan et al., 2015), although precise correlations require further analyses.

**Figure 3.**
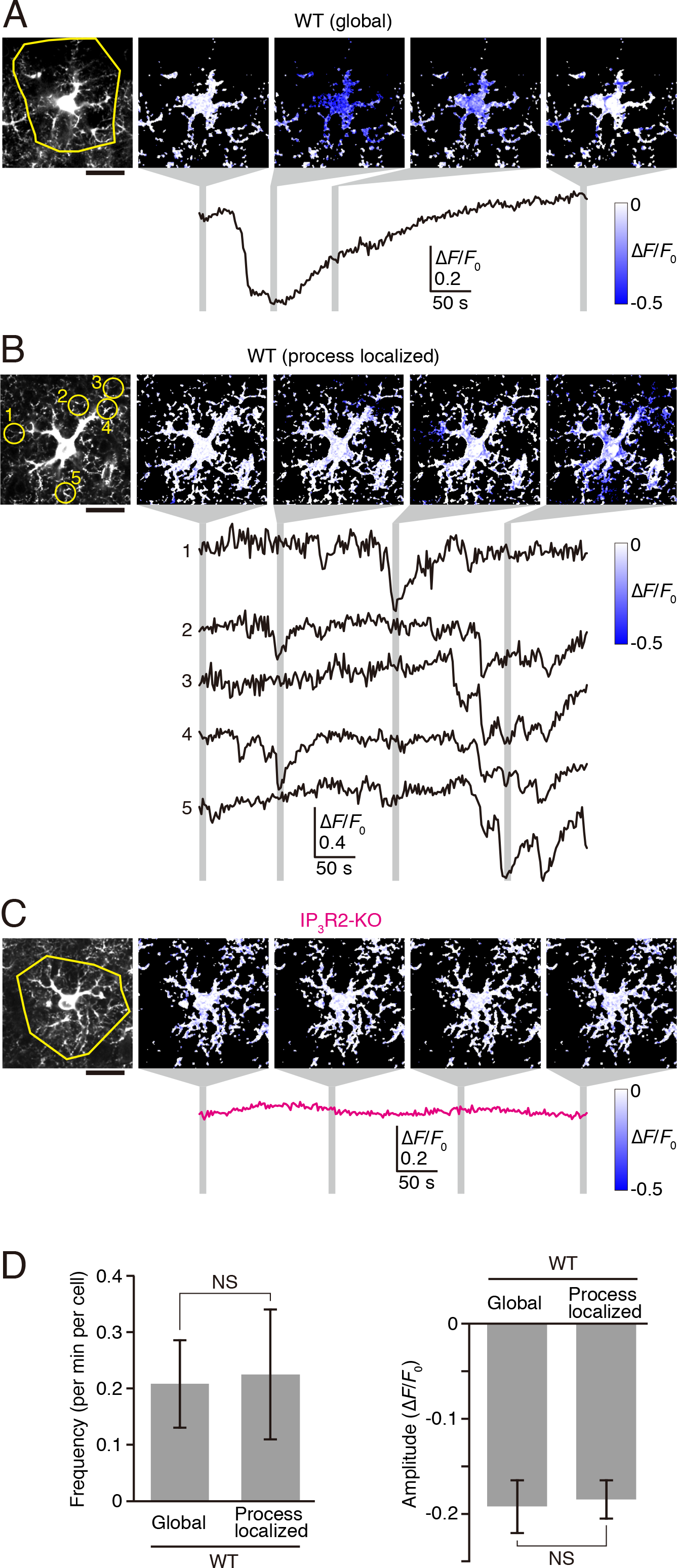
Spontaneous ER Ca^2+^ release in WT cortical astrocytes. (A and B) Representative spontaneous G-CEPIA1*er* responses that emerged in WT astrocytes either synchronously throughout the astrocyte (global response, A) or in confined regions of the processes (process-localized response, B). The yellow circles in the leftmost images indicate the region of interest for the Δ*F*/*F*_0_ traces. Pseudocolor images indicate the average of five consecutive Δ*F*/*F*_0_ images within the time zones indicated by gray vertical bars. Scale bar, 20 μm. (C) Representative G-CEPIA1*er* responses in IP_3_R2-KO astrocytes without external stimulation. Data were shown as in A and B. (D) Summary of frequencies (left) and amplitudes (right) of global and process-localized responses in WT astrocytes. *n* = 25 global responses and 27 process-localized responses in 15 cells (mean ± s.e.m.). No significant difference was detected between global and process-localized responses (*P* = 0.905 for frequency, *P* = 0.831 for amplitude, unpaired Student’s *t*-test).

### IP_3_R2-independent Ca^2+^ release in hippocampal astrocytes

Regional differences in the properties of Ca^2+^ signaling in astrocytes have been reported (Chai et al., 2017). To investigate whether IP_3_R2-independent Ca^2+^ release could be observed in astrocytes of brain regions other than the cortex, we visualized Ca^2+^ release from the ER in hippocampal astrocytes. G-CEPIA1*er* was expressed in hippocampal astrocytes using the AAV. G-CEPIA1*er*-expressing astrocytes in the CA1 stratum radiatum of acute hippocampal slice preparations were imaged by two-photon microscopy. Application of NE induced large G-CEPIA1*er* responses in WT astrocytes (Fig. 4A and B), whereas attenuated G-CEPIA1*er* responses were observed in IP_3_R2-KO astrocytes (Fig. 4A and B). Therefore, IP_3_R2-independent Ca^2+^ release was commonly observed in cortical and hippocampal astrocytes of IP_3_R2-KO mice. “Global” and “process localized” spontaneous G-CEPIA1*er* responses were observed in WT astrocytes, but not in IP_3_R2-KO astrocytes, which was consistent with the results in cortical astrocytes (Fig. 4).

**Figure 4.**
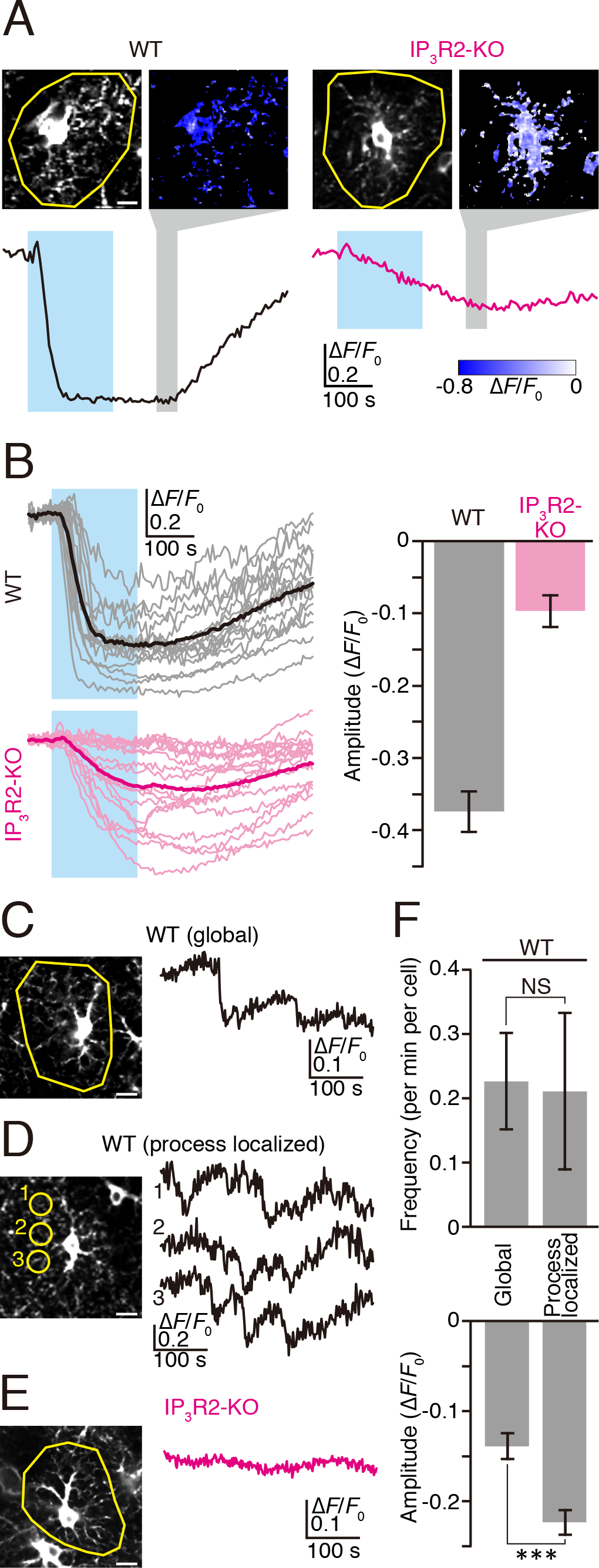
IP_3_R2-independent Ca^2+^ release in hippocampal astrocytes. (A) Representative G-CEPIA1*er* responses upon application of NE (10 μM, cyan bars) to WT and IP_3_R2-KO astrocytes in hippocampal slices. Yellow circles indicate the region of interest for the Δ*F*/*F*_0_ traces (black trace for WT and magenta trace for IP_3_R2-KO). Pseudocolor images indicate the average of ten consecutive Δ*F*/*F*_0_ images within the time zones indicated by gray vertical bars. Scale bar, 10 μm. (B) Left: G-CEPIA1*er* responses upon application of NE (10 μM, cyan bars) to WT (black) and IP_3_R2-KO (magenta) astrocytes. Individual Δ*F*/*F*_0_ traces (pale color) and average traces (deep color) are indicated. Right: Summary of the response amplitudes of Δ*F*/*F*_0_ traces (*n* = 16 cells for WT and 18 cells for IP_3_R2-KO, mean ± s.e.m.). (C and D) Representative spontaneous G-CEPIA1*er* responses that emerged in WT hippocampal astrocytes either synchronously throughout the astrocyte (global response, C) or in confined regions of the processes (process-localized response, D). The yellow circles in the image indicate the region of interest for the Δ*F*/*F*_0_ traces. Scale bar, 10 μm. (E) Representative G-CEPIA1*er* responses in IP_3_R2-KO hippocampal astrocytes without external stimulation. Data were shown as in C and D. (F) Summary of frequencies and amplitudes of global and process-localized responses in hippocampal WT astrocytes. *n* = 29 global responses and 27 process-localized responses in 16 cells (mean ± s.e.m.). No significant difference was detected in frequency of global and process-localized responses (*P* = 0.914, unpaired Student’s *t*-test). Amplitude of process-localized responses was significantly larger than global responses (\*\*\**P* < 0.001, unpaired Student’s *t*-test).

### Minimal Ca^2+^ transients in the cytosol mediated by IP_3_R2-independent Ca^2+^ release

Having found significant IP_3_R2-independent Ca^2+^ release in IP_3_R2-KO astrocytes, we decided to reinvestigate changes in the cytosolic Ca^2+^ concentration ([Ca^2+^]_Cyt_) in response to NE application (Fig. 4). Cytosolic Ca^2+^ indicator GCaMP6f (Chen et al., 2013) was expressed in cortical astrocytes using an AAV, showing a diffuse distribution throughout the cytosol (Fig. 4A). In WT astrocytes, application of NE induced large [Ca^2+^]_Cyt_ responses that were clearly detected by Δ*F*/*F*_0_ traces of GCaMP6f averaged within the entire cell (Fig. 4A). Responses were abolished after treatment with CPA, indicating a critical contribution of ER Ca^2+^ release to NE-induced [Ca^2+^]_Cyt_ transients in WT astrocytes (Fig. 4B and C). On the other hand, NE induced very little [Ca^2+^]_Cyt_ responses in IP_3_R2-KO astrocytes (Fig. 4A). Most IP_3_R2-KO astrocytes showed no or minimal responses, and only very few IP_3_R2-KO astrocytes showed clear but relatively small responses (Fig. 4B). The [Ca^2+^]_Cyt_ responses in IP_3_R2-KO astrocytes were also abolished after treatment with CPA (Fig. 4B and C). These results indicate that IP_3_R2-independent Ca^2+^ release, which was detected by G-CEPIA1*er*, did not always increase [Ca^2+^]_Cyt_ to a level detectable by GCaMP6f.

### Robust Ca^2+^ transients in mitochondria mediated by IP_3_R2-independent Ca^2+^ release

The results so far indicate that IP_3_R2-independent Ca^2+^ release is clearly present in astrocytes, but that it induces only very little cytosolic Ca^2+^ signals. Does it mean IP_3_R2-independent Ca^2+^ release has no physiological role? It has been shown that the ER and mitochondria form close contacts where Ca^2+^ is efficiently transferred from the ER to mitochondria without a global increase in [Ca^2+^]_Cyt_ (Rizzuto et al., 1998; de Brito & Scorrano, 2010; Hirabayashi et al., 2017). This mitochondrial Ca^2+^ signaling regulates various physiological and pathophysiological functions of mitochondria (Rizzuto, De Stefani, Raffaello, & Mammucari, 2012). We, therefore, examined whether IP_3_R2-independent Ca^2+^ release would contribute to the ER-mitochondrial coupling. To study changes in the Ca^2+^ concentration within mitochondria ([Ca^2+^]_Mito_), we used CEPIA2*mt*, a Ca^2+^ indicator for the mitochondrial matrix (Suzuki et al., 2014) (Fig. 5). AAV-mediated expression of CEPIA2*mt* in cortical astrocytes resulted in a punctate distribution of fluorescence, which was consistent with the morphology of mitochondria (Fig. 5A). Application of NE to release Ca^2+^ from the ER induced an increase in [Ca^2+^]_Mito_ of WT astrocytes (Fig. 5B). Although a subpopulation of WT astrocytes showed no or very little responses (Fig. 5C), such intercellular heterogeneity of mitochondrial Ca^2+^ signaling has been similarly observed in HeLa cells (Suzuki et al., 2014). The NA-induced [Ca^2+^]_Mito_ response was abolished after treatment with CPA to deplete the ER (Fig. 5C and D). These results indicate that Ca^2+^ enters mitochondria following Ca^2+^ release from the ER in WT astrocytes.

**Figure 5.**
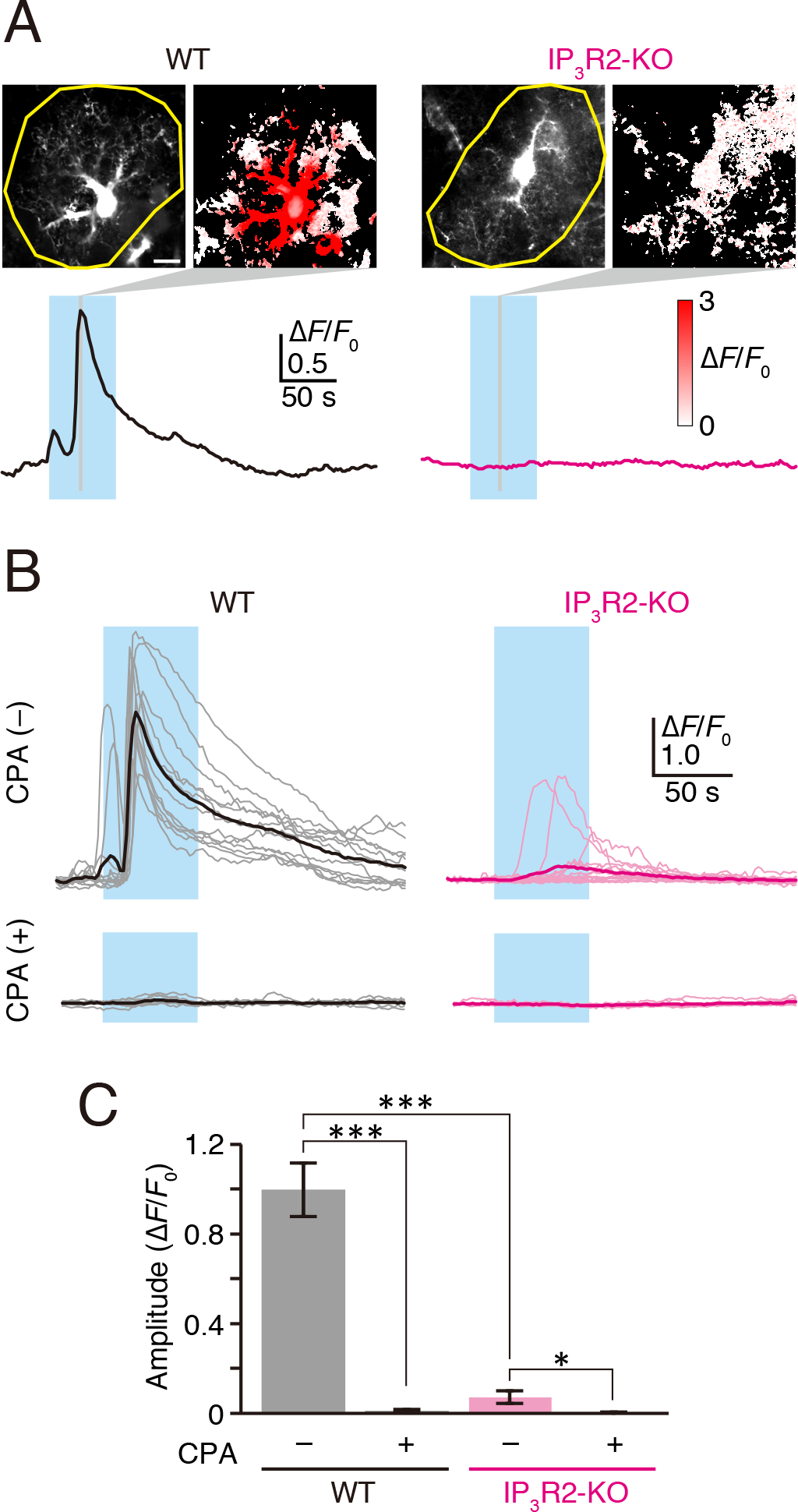
Cytosolic Ca^2+^ transients induced by IP_3_R2-independent Ca^2+^ release. (A) Representative GCaMP6f responses upon application of NE (10 μM, cyan bars) to WT and IP_3_R2-KO astrocytes. The yellow circles indicate the region of interest for the Δ*F*/*F*_0_ traces (black trace for WT and magenta trace for IP_3_R2-KO). Pseudocolor images indicate Δ*F*/*F*_0_ at the time points indicated by gray vertical bars in the Δ*F*/*F*_0_ traces. Fluorescence increase in the soma of WT astrocytes was underestimated due to saturation of the dynamic range of photodetector. Scale bar, 10 μm. (B) GCaMP6f responses upon application of NE (10 μM, cyan bars) to WT (black) and IP_3_R2-KO (magenta) astrocytes in the control condition [CPA (−)] or after treatment with CPA [CPA (+)]. A whole astrocyte was selected as the region of interest to measure Δ*F*/*F*_0_. Individual Δ*F*/*F*_0_ traces (pale color) and average traces (deep color) are indicated. *n* = 13 cells for WT [CPA (−)], 10 cells for WT [CPA (+)], 16 cells for IP_3_R2-KO [CPA (−)], and 10 cells for IP_3_R2-KO [CPA (+)]. (C) Summary of the response amplitudes of Δ*F*/*F*_0_ traces shown in (B) (mean ± s.e.m.). Asterisks indicate the statistical significance. \**P* < 0.05, \*\*\**P* < 0.001, unpaired Student’s *t*-test.

**Figure 6.**
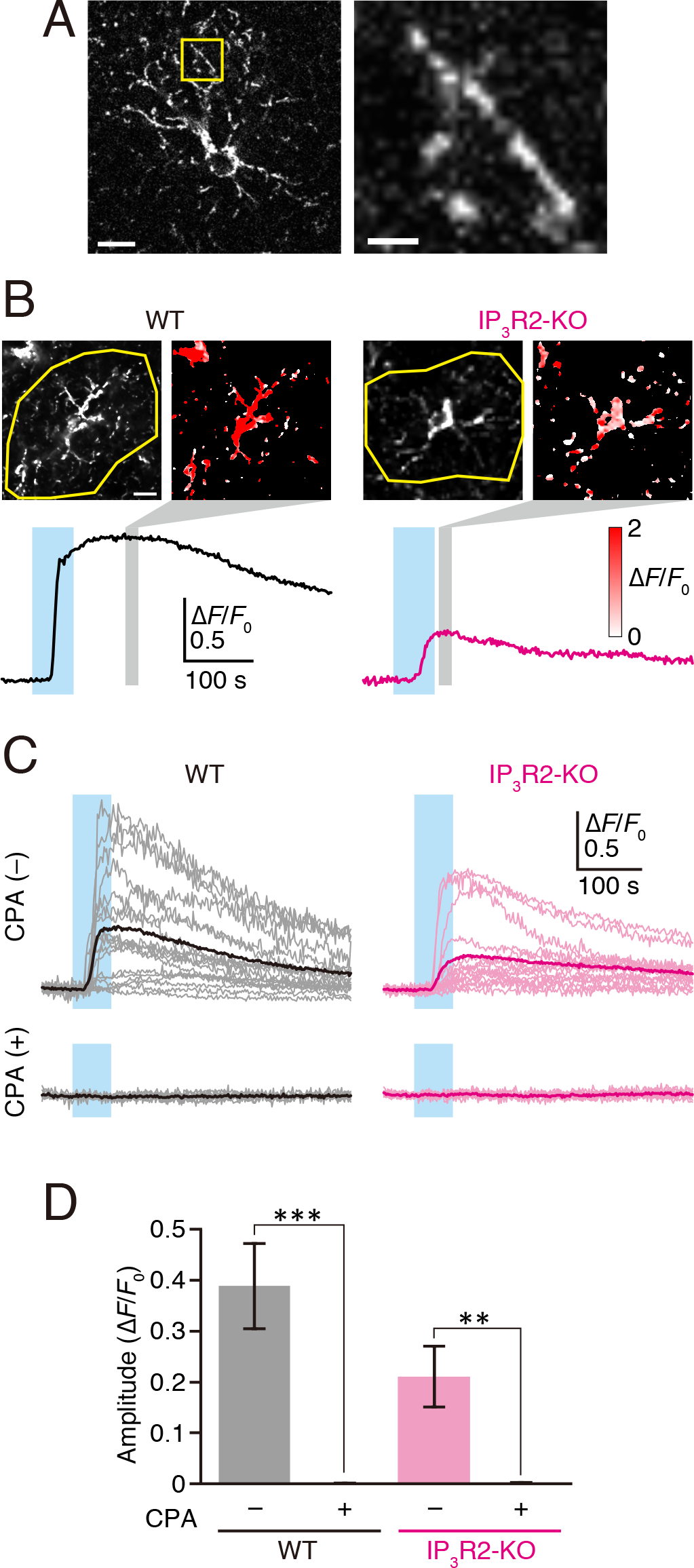
Mitochondrial Ca^2+^ transients induced by IP_3_R2-independent Ca^2+^ release. (A) CEPIA2*mt*-expressing astrocyte. The right image is the magnification of the region indicated by the yellow box in the left image. Scale bars, 10 μm (left) and 2 μm (right). (B) Representative CEPIA2*mt* responses upon application of NE (10 μM, cyan bars) to WT and IP_3_R2-KO astrocytes. The yellow circles indicate the region of interest for the Δ*F*/*F*_0_ traces (black trace for WT and magenta trace for IP_3_R2-KO). Pseudocolor images indicate the average of ten consecutive Δ*F*/*F*_0_ images within the time zones indicated by gray vertical bars. Scale bar, 10 μm. (C) CEPIA2*mt* responses upon application of NE (10 μM, cyan bars) to WT (black) and IP_3_R2-KO (magenta) astrocytes in the control condition [CPA (−)] or after treatment with CPA [CPA (+)]. A whole astrocyte was selected as the region of interest to measure Δ*F*/*F*_0_. Individual Δ*F*/*F*_0_ traces (pale color) and average traces (deep color) are shown. *n* = 18 cells for WT [CPA (−)], 10 cells for WT [CPA (+)], 16 cells for IP_3_R2-KO [CPA (−)], and 12 cells for IP_3_R2-KO [CPA (+)]. (D) Summary of the response amplitudes of Δ*F*/*F*_0_ traces shown in (B) (mean ± s.e.m.), indicating significant CEPIA2*mt* responses in IP_3_R2-KO astrocytes. Absence of response after CPA indicates the requirement of ER Ca^2+^ release to CEPIA2*mt* responses. Asterisks indicate the statistical significance of the responses in CPA (−) condition versus CPA (+) condition in each genotype. \*\**P* < 0.01, \*\*\**P* < 0.001, unpaired Student’s *t*-test. Difference between WT [CPA (−)] and IP_3_R2-KO [CPA (−)] responses was not statistically significant (*P* = 0.0553, unpaired Student’s *t*-test).

Of note, NE also induced [Ca^2+^]_Mito_ responses in IP_3_R2-KO astrocytes (Fig. 5A). Similar to WT astrocytes, a subpopulation of IP_3_R2-KO astrocytes showed no or very little responses. Compared with the minimal [Ca^2+^]_Cyt_ responses, [Ca^2+^]_Mito_ showed significant responses in IP_3_R2-KO astrocytes upon NE stimulation. The [Ca^2+^]_Mito_ responses in IP_3_R2-KO astrocytes were also abolished after treatment with CPA (Fig. 5B and C). These results indicate that IP_3_R2-independent Ca^2+^ release efficiently induces an increase in [Ca^2+^]_Mito_.

## Discussion

The results of the present study provide clear evidence that Ca^2+^ is released from the ER in IP_3_R2-KO astrocytes upon agonist stimulation. Although the IP_3_R2-independent Ca^2+^ release from the ER was ineffective in increasing the cytosolic Ca^2+^ concentration in IP_3_R2-KO astrocytes, it was effective to increase the Ca^2+^ concentration within mitochondria that make close contacts with the ER. These results provide new insights into the functional significance of ER Ca^2+^ release in astrocytes. The controversy derived from the assumption that ER Ca^2+^ release is abolished in IP_3_R2-KO astrocytes should be re-evaluated considering the presence of IP_3_R2-independent Ca^2+^ release.

### Highly sensitive and specific detection of ER Ca^2+^ release using G-CEPIA1*er*

Although IP_3_R2-independent Ca^2+^ release from the ER in IP_3_R2-KO astrocytes was detectable by G-CEPIA1*er* responses, the resulting changes in [Ca^2+^]_Cyt_ were only barely detected by GCaMP6f responses. The difficulty in detecting the [Ca^2+^]_Cyt_ response is consistent with previous reports that failed to detect evoked [Ca^2+^]_Cyt_ responses in IP_3_R2-KO astrocytes (Petravicz, Fiacco, & McCarthy, 2008; Agulhon, Fiacco, & McCarthy, 2010; Agulhon et al., 2013; Nizar et al., 2013; Takata et al., 2013; Petravicz, Boyt, & McCarthy, 2014). These results suggest that the amount of Ca^2+^ release from the ER sufficient to change free [Ca^2+^]_ER_ might be insufficient to change free [Ca^2+^]_Cyt_. There are a couple of potential reasons for the puzzling observation. First, the volume occupied by the smooth ER (*V*_ER_) is about 10% of that of the cytoplasm (*V*_Cyt_) (Paumgartner, Losa, & Weibel, 1981). Therefore, Ca^2+^ released from the ER is 10-fold diluted in the cytosol. Another possible reason is the difference in the Ca^2+^ buffering capacity *k* (a change in the total Ca^2+^ concentration in a subcellular compartment divided by the corresponding change in free Ca^2+^ concentration in the compartment) between the ER and cytosol. In many cell types, cytosolic *k* (*k*_Cyt_) is estimated to be 10–2,000 due mainly to the presence of high concentrations of Ca^2+^ binding proteins in the cytosol (Allbritton, Meyer, & Stryer, 1992; Neher & Augustine, 1992; Neher, 1995; Mogami et al., 1999). However, *k* of the ER (*k*_ER_) is estimated to be two orders of magnitude less than *k*_Cyt_ in pancreatic acinar cells (Mogami et al., 1999). If *k*_ER_ is similarly lower than *k*_Cyt_ in astrocytes, this would also reduce the Ca^2+^ release-induced change in [Ca^2+^]_Cyt_. We thus estimated the dilution and buffering effects in astrocytes. We previously measured both [Ca^2+^]_Cyt_ and [Ca^2+^]_ER_ in astrocytes stimulated with 30 μM ATP (Suzuki et al., 2014). Since the peak changes in [Ca^2+^]_Cyt_ and [Ca^2+^]_ER_ were observed almost simultaneously, we assumed that the change in the total Ca^2+^ concentration in the cytosol can be approximated by that of ER Ca^2+^ concentration as follows:

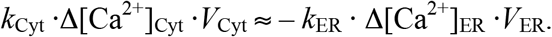

Δ[Ca^2+^]_Cyt_ was similar to those of HeLa, HEK, and BHK cells, and most likely to be of the order of 1 μM, while Δ[Ca^2+^]_ER_ estimated by the ratiometric ER Ca^2+^ indicator GEM-CEPIA1*er* was about –260 μM (Suzuki et al., 2014). Thus, the product of buffering and volume factors can be described as follows:

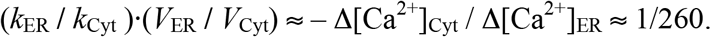

This indicates that Ca^2+^ release from the ER is highly buffered and diluted in the cytosol. In addition to the presence of Ca^2+^ binding proteins, effects of the efflux of Ca^2+^ to the extracellular space and influx to other intracellular organelles such as mitochondria would partly contribute to the estimated buffering/dilution factor. Thus, free Ca^2+^ concentration changes in the cytosol are only about 0.4% of those in the ER. Therefore, G-CEPIA1*er* is expected to report ER Ca^2+^ release with much higher sensitivity than cytosolic Ca^2+^ indicators. Furthermore, G-CEPIA1*er* can specifically detect ER Ca^2+^ release, whereas cytosolic Ca^2+^ indicators do not specify the source of Ca^2+^.

### Contribution of IP_3_R2-independent Ca^2+^ release to Ca^2+^ signaling in IP_3_R2-KO astrocytes

When the extent of the NE-induced increase in Ca^2+^ concentration was compared between WT and IP_3_R2-KO astrocytes, there was a much higher Ca^2+^ increase in mitochondria than in the cytoplasm of IP_3_R2-KO astrocytes (Figs. 4 and 5). These results suggest an IP_3_R2-independent mechanism that allows privileged transfer of Ca^2+^ from the ER to mitochondria. Indeed, it has been shown that the ER and mitochondria make close contacts (10–30 nm in distance), and that such contact sites are crucial for efficient Ca^2+^ transfer (Rizzuto et al., 1998; de Brito & Scorrano, 2010; Hirabayashi et al., 2017). This finding suggests that IP_3_R2-independent Ca^2+^ release is capable of increasing [Ca^2+^]_Cyt_ within close proximity to the ER membrane, generating Ca^2+^ nanodomains around the ER (Rizzuto & Pozzan, 2006). These Ca^2+^ nanodomains generated within the ER-mitochondria contact sites should mediate efficient Ca^2+^ transfer between these two organelles (Rizzuto, De Stefani, Raffaello, & Mammucari, 2012). Other functional significances of IP_3_R2-independent Ca^2+^ nanodomains around the ER require further clarification. Thus, in addition to extracellular and mitochondrial sources of Ca^2+^ (Srinivasan et al., 2015; Agarwal et al., 2017), IP_3_R2-independent Ca^2+^ release from the ER may contribute to Ca^2+^ signaling in IP_3_R2-KO astrocytes.

### Ca^2+^ channels that mediate IP_3_R2-independent Ca^2+^ release

Although IP_3_R2 is the major ER Ca^2+^ release channel in astrocytes, expression of IP_3_R1 and/or IP_3_R3 in astrocytes has been indicated by transcriptome analyses (Cahoy et al., 2008; Zhang et al., 2014; Chai et al., 2017) and immunohistochemical studies (Yamamoto-Hino et al., 1995; Sharp et al., 1999). Furthermore, comparison between IP_3_R2-KO mice and IP_3_R2/IP_3_R3-double KO mice indicate a functional contribution of IP_3_R3 to astrocytic Ca^2+^ signaling, albeit small (Tamamushi et al., 2012; Sherwood et al., 2017). These previous studies suggest the presence of IP_3_-induced Ca^2+^ release via IP_3_R1 and/or IP_3_R3 in IP_3_R2-KO astrocytes. Indeed, crucial roles of IP_3_R1 and IP_3_R3 for the ER-mitochondria coupling have been reported. IP_3_R1 has been shown to be one of the components that physically link the ER to mitochondria (Szabadkai et al., 2006). IP_3_R3 is considered as a major subtype that mediates Ca^2+^ transfer from the ER to mitochondria (Mendes et al., 2005; Hayashi & Su, 2007; Cárdenas et al., 2010). Therefore, the involvement of IP_3_R1 and/or IP_3_R3 in mitochondrial Ca^2+^ signaling in IP_3_R2-KO astrocytes is now being addressed.

In addition, expression of ryanodine receptor type 3 (RyR3) in astrocytes has been reported (Matyash et al., 2002; Chai et al., 2017). The functional significance of RyR3 was reported in both astrocytic Ca^2+^ signaling and motility (Matyash et al., 2002). Therefore, it also appears possible that Ca^2+^-induced Ca^2+^ release via RyR3 may enhance Ca^2+^ release from the ER via IP_3_R1 and/or IP_3_R3.

### Reinterpretation of astrocytic functions in IP_3_R2-KO mice

It has been assumed that Ca^2+^ release from the ER is absent in IP_3_R2-KO astrocytes. Thus, the absence of functional deficits in IP_3_R2-KO mice (Petravicz, Fiacco, & McCarthy, 2008; Agulhon, Fiacco, & McCarthy, 2010; Agulhon et al., 2013; Nizar et al., 2013; Takata et al., 2013; Bonder & McCarthy, 2014; Petravicz, Boyt, & McCarthy, 2014) has been considered to indicate the absence of functional roles of intracellular Ca^2+^ release in astrocytes. The presence of IP_3_R2-independent Ca^2+^ release may necessitate reinterpretation of such results.

Ca^2+^ release from the ER may be inhibited by expressing the IP_3_ hydrolyzing enzyme, IP_3_ 5-phosphatase (5ppase) (Kanemaru, Okubo, Hirose, & Iino, 2007; Mashimo et al., 2010; Foley et al., 2017). Indeed, exogenous expression of 5ppase effectively suppresses IP_3_ production following IP_3_R-dependent Ca^2+^ release in astrocytes (Kanemaru, Okubo, Hirose, & Iino, 2007; Mashimo et al., 2010). Because 5ppase is expected to suppress IP_3_R-induced Ca^2+^ release regardless of IP_3_R subtypes, the phenotypic difference between 5ppase-expressing mice and IP_3_R2-KO mice may clarify the role of IP_3_R2-independent Ca^2+^ release. In fact, a recent study has indicated that astrocytic expression of 5ppase disrupts sleep (Foley et al., 2017), whereas sleep disruption was not observed in IP_3_R2-KO mice (Cao et al., 2013). Although there remains the possibility of developmental adaptation or compensatory mechanisms in these mice, the results suggest that residual Ca^2+^ signaling, including IP_3_R2-independent Ca^2+^ release in IP_3_R2-KO mice, accounts for the absence of functional deficits in IP_3_R2-KO mice.

In conclusion, the presence of IP_3_R2-independent Ca^2+^ release as well as Ca^2+^ signaling generated by sources other than the ER limit the use of IP_3_R2-KO mice to assess the role of Ca^2+^ signaling in astrocytic functions when functional deficits are not observed in the mutant mice. This notion may help to resolve the long-lasting controversy arising from the assumption that Ca^2+^ signaling is abolished in IP_3_R2-KO astrocytes. Furthermore, the present results facilitate the study of the functional roles of Ca^2+^ signaling in close proximity to the ER in astrocytes.

## Acknowledgements

We thank Y. Kawashima for technical assistance and Dr. B. Khakh (University of California at San Francisco) for providing the pZac2.1-gfaABC1D-Lck-GCaMP3 plasmid. This work was supported by the Japan Society for the Promotion of Science (JSPS) KAKENHI [Grant Numbers JP16K08543 (Y.O.), JP15H05648 (K.K.), and JP25117002 and JP25221304 (M.I.)] and grants from the Tokyo Society of Medical Sciences (Y.O.).

## Notes

**Conflict of Interest Statement** The authors declare no competing financial interests.

## References

Agarwal, A., Wu, P.-H., Hughes, E. G., Fukaya, M., Tischfield, M. A., Langseth, A. J., … Bergles, D. E. (2017). Transient opening of the mitochondrial permeability transition pore induces microdomain calcium transients in astrocyte processes. Neuron 93, 587–605.

Agulhon, C., Boyt, K. M., Xie, A. X., Friocourt, F., Roth, B. L., & McCarthy, K. D. (2013). Modulation of the autonomic nervous system and behaviour by acute glial cell G_q_ protein-coupled receptor activation in vivo. The Journal of Physiology 591, 5599–5609.

Agulhon, C., Fiacco, T. A., & McCarthy, K. D. (2010). Hippocampal short- and long-term plasticity are not modulated by astrocyte Ca^2+^ signaling. Science 327, 1250–1254.

Allbritton, N., Meyer, T., & Stryer, L. (1992). Range of messenger action of calcium ion and inositol 1,4,5-trisphosphate. Science 258, 1812–1815.

Barres, B. A. (2008). The mystery and magic of glia: a perspective on their roles in health and disease. Neuron 60, 430–440.

Bazargani, N., & Attwell, D. (2016). Astrocyte calcium signaling: the third wave. Nature Neuroscience 19, 182–189.

Bonder, D. E., & McCarthy, K. D. (2014). Astrocytic G_q_-GPCR-linked IP_3_R-dependent Ca^2+^ signaling does not mediate neurovascular coupling in mouse visual cortex in vivo. Journal of Neuroscience 34, 13139–13150.

de Brito, O. M., & Scorrano, L. (2010). An intimate liaison: spatial organization of the endoplasmic reticulum-mitochondria relationship. The EMBO journal 29, 2715–23.

Cahoy, J. D., Emery, B., Kaushal, A., Foo, L. C., Zamanian, J. L., Christopherson, K. S., … Barres, B. A. (2008). A transcriptome database for astrocytes, neurons, and oligodendrocytes: a new resource for understanding brain development and function. Journal of Neuroscience 28, 264–278.

Cao, X., Li, L.-P., Wang, Q., Wu, Q., Hu, H.-H., Zhang, M., … Gao, T.-M. (2013). Astrocyte-derived ATP modulates depressive-like behaviors. Nature Medicine 19, 773–777.

Cárdenas, C., Miller, R. A., Smith, I., Bui, T., Molgó, J., Müller, M., … Foskett, J. K. (2010). Essential regulation of cell bioenergetics by constitutive InsP_3_ receptor Ca^2+^ transfer to mitochondria. Cell 142, 270–283.

Carreras-Sureda, A., Pihán, P., & Hetz, C. (2018). Calcium signaling at the endoplasmic reticulum: fine-tuning stress responses. Cell Calcium 70, 24–31.

Chai, H., Diaz-Castro, B., Shigetomi, E., Monte, E., Octeau, J. C., Yu, X., … Khakh, B. S. (2017). Neural circuit-specialized astrocytes: transcriptomic, proteomic, morphological, and functional evidence. Neuron 95, 531–549.

Chen, N., Sugihara, H., Sharma, J., Perea, G., Petravicz, J., Le, C., & Sur, M. (2012). Nucleus basalis-enabled stimulus-specific plasticity in the visual cortex is mediated by astrocytes. Proceedings of the National Academy of Sciences 109, E2832–E2841.

Chen, T.-W., Wardill, T. J., Sun, Y., Pulver, S. R., Renninger, S. L., Baohan, A., … Kim, D. S. (2013). Ultrasensitive fluorescent proteins for imaging neuronal activity. Nature 499, 295–300.

Ding, F., O’Donnell, J., Thrane, A. S., Zeppenfeld, D., Kang, H., Xie, L., … Nedergaard, M. (2013). α1-Adrenergic receptors mediate coordinated Ca^2+^ signaling of cortical astrocytes in awake, behaving mice. Cell Calcium 54, 387–394.

Dong, Q., He, J., & Chai, Z. (2013). Astrocytic Ca^2+^ waves mediate activation of extrasynaptic NMDA receptors in hippocampal neurons to aggravate brain damage during ischemia. Neurobiology of Disease 58, 68–75.

Edwards, F. A., Konnerth, A., Sakmann, B., & Takahashi, T. (1989). A thin slice preparation for patch clamp recordings from neurones of the mammalian central nervous system. Pflügers Archiv: European journal of physiology 414, 600–12.

Fiacco, T. A., & McCarthy, K. D. (2018). Multiple lines of evidence indicate that gliotransmission does not occur under physiological conditions. The Journal of neuroscience: the official journal of the Society for Neuroscience 38, 3–13.

Foley, J., Blutstein, T., Lee, S., Erneux, C., Halassa, M. M., & Haydon, P. (2017). Astrocytic IP_3_/Ca^2+^ signaling modulates theta rhythm and REM sleep. Frontiers in neural circuits 11, 3.

Hamilton, N. B., & Attwell, D. (2010). Do astrocytes really exocytose neurotransmitters? Nature Reviews Neuroscience 11, 227–238.

Hayashi, T., & Su, T.-P. (2007). Sigma-1 receptor chaperones at the ER- mitochondrion interface regulate Ca^2+^ signaling and cell survival. Cell 131, 596–610.

Hirabayashi, Y., Kwon, S.-K., Paek, H., Pernice, W. M., Paul, M. A., Lee, J., … Polleux, F. (2017). ER-mitochondria tethering by PDZD8 regulates Ca^2+^ dynamics in mammalian neurons. Science (New York, NY) 358, 623–630.

Holtzclaw, L. A., Pandhit, S., Bare, D. J., Mignery, G. A., & Russell, J. T. (2002). Astrocytes in adult rat brain express type 2 inositol 1,4,5-trisphosphate receptors. Glia 39, 69–84.

Kanemaru, K., Kubota, J., Sekiya, H., Hirose, K., Okubo, Y., & Iino, M. (2013). Calcium-dependent N-cadherin up-regulation mediates reactive astrogliosis and neuroprotection after brain injury. Proceedings of the National Academy of Sciences 110, 11612–11617.

Kanemaru, K., Okubo, Y., Hirose, K., & Iino, M. (2007). Regulation of neurite growth by spontaneous Ca^2+^ oscillations in astrocytes. The Journal of neuroscience: the official journal of the Society for Neuroscience 27, 8957–8966.

Kanemaru, K., Sekiya, H., Xu, M., Satoh, K., Kitajima, N., Yoshida, K., … Tanaka, K. F. (2014). In vivo visualization of subtle, transient, and local activity of astrocytes using an ultrasensitive Ca^2+^ indicator. Cell Reports 8, 311–318.

Kim, S. K., Hayashi, H., Ishikawa, T., Shibata, K., Shigetomi, E., Shinozaki, Y., … Nabekura, J. (2016). Cortical astrocytes rewire somatosensory cortical circuits for peripheral neuropathic pain. The Journal of Clinical Investigation 126, 1983–1997.

Li, H., Xie, Y., Zhang, N., Yu, Y., Zhang, Q., & Ding, S. (2015). Disruption of IP_3_R2-mediated Ca^2+^ signaling pathway in astrocytes ameliorates neuronal death and brain damage while reducing behavioral deficits after focal ischemic stroke. Cell Calcium 58, 565–576.

Li, X., Zima, A. V., Sheikh, F., Blatter, L. A., & Chen, J. (2005). Endothelin-1–induced arrhythmogenic Ca^2+^ signaling is abolished in atrial myocytes of inositol-1,4,5-trisphosphate (IP_3_)–receptor type 2–deficient mice. Circulation Research 96, 1274–81.

Mashimo, M., Okubo, Y., Yamazawa, T., Yamasaki, M., Watanabe, M., Murayama, T., & Iino, M. (2010). Inositol 1,4,5-trisphosphate signaling maintains the activity of glutamate uptake in Bergmann glia. European Journal of Neuroscience 32, 1668–1677.

Matyash, M., Matyash, V., Nolte, C., Sorrentino, V., & Kettenmann, H. (2002). Requirement of functional ryanodine receptor type 3 for astrocyte migration. The FASEB Journal 16, 84–86.

Mendes, C. C. P., Gomes, D. A., Thompson, M., Souto, N. C., Goes, T. S., Goes, A. M., … Leite, M. F. (2005). The type III inositol 1,4,5-trisphosphate receptor preferentially transmits apoptotic Ca^2+^ signals into mitochondria. The Journal of biological chemistry 280, 40892–900.

Mogami, H., Gardner, J., Gerasimenko, O., Camello, P., Petersen, O., & Tepikin, A. (1999). Calcium binding capacity of the cytosol and endoplasmic reticulum of mouse pancreatic acinar cells. The Journal of physiology 518, 463–7.

Monai, H., Ohkura, M., Tanaka, M., Oe, Y., Konno, A., Hirai, H., … Hirase, H. (2016). Calcium imaging reveals glial involvement in transcranial direct current stimulation-induced plasticity in mouse brain. Nature Communications 7, 11100.

Navarrete, M., Perea, G., de Sevilla, D. F., Gómez-Gonzalo, M., Núñez, A., Martín, E. D., & Araque, A. (2012). Astrocytes mediate in vivo cholinergic-induced synaptic plasticity. PLoS Biol 10, e1001259.

Neher, E. (1995). The use of fura-2 for estimating Ca buffers and Ca fluxes. Neuropharmacology 34, 1423–1442.

Neher, E., & Augustine, G. J. (1992). Calcium gradients and buffers in bovine chromaffin cells. The Journal of physiology 450, 273–301.

Nizar, K., Uhlirova, H., Tian, P., Saisan, P. A., Cheng, Q., Reznichenko, L., … Devor, A. (2013). In vivo stimulus-induced vasodilation occurs without IP_3_ receptor activation and may precede astrocytic calcium increase. Journal of Neuroscience 33, 8411–8422.

Okubo, Y., Suzuki, J., Kanemaru, K., Nakamura, N., Shibata, T., & Iino, M. (2015). Visualization of Ca^2+^ filling mechanisms upon synaptic inputs in the endoplasmic reticulum of cerebellar Purkinje cells. Journal of Neuroscience 35, 15837–15846.

Padmashri, R., Suresh, A., Boska, M. D., & Dunaevsky, A. (2015). Motor-skill learning is dependent on astrocytic activity. Neural Plasticity 2015, 938023.

Patrushev, I., Gavrilov, N., Turlapov, V., & Semyanov, A. (2013). Subcellular location of astrocytic calcium stores favors extrasynaptic neuron–astrocyte communication. Cell Calcium 54, 343–349.

Paukert, M., Agarwal, A., Cha, J., Doze, V. A., Kang, J. U., & Bergles, D. E. (2014). Norepinephrine controls astroglial responsiveness to local circuit activity. Neuron 82, 1263–1270.

Paumgartner, D., Losa, G., & Weibel, E. R. (1981). Resolution effect on the stereological estimation of surface and volume and its interpretation in terms of fractal dimensions. Journal of Microscopy 121, 51–63.

Perez-Alvarez, A., Navarrete, M., Covelo, A., Martin, E. D., & Araque, A. (2014). Structural and functional plasticity of astrocyte processes and dendritic spine interactions. Journal of Neuroscience 34, 12738–12744.

Petravicz, J., Boyt, K. M., & McCarthy, K. D. (2014). Astrocyte IP_3_R2-dependent Ca^2+^ signaling is not a major modulator of neuronal pathways governing behavior. Frontiers in Behavioral Neuroscience 8, 384.

Petravicz, J., Fiacco, T. A., & McCarthy, K. D. (2008). Loss of IP_3_ receptor-dependent Ca^2+^ increases in hippocampal astrocytes does not affect baseline CA1 pyramidal neuron synaptic activity. The Journal of neuroscience: the official journal of the Society for Neuroscience 28, 4967–73.

Rakers, C., & Petzold, G. C. (2016). Astrocytic calcium release mediates peri-infarct depolarizations in a rodent stroke model. Journal of Clinical Investigation 127, 511–516.

Rizzuto, R., Pinton, P., Carrington, W., Fay, F. S., Fogarty, K. E., Lifshitz, L. M., … Pozzan, T. (1998). Close contacts with the endoplasmic reticulum as determinants of mitochondrial Ca^2+^ responses. Science (New York, NY) 280, 1763–1766.

Rizzuto, R., & Pozzan, T. (2006). Microdomains of intracellular Ca^2+^: molecular determinants and functional consequences. Physiological Reviews 86, 369–408.

Rizzuto, R., De Stefani, D., Raffaello, A., & Mammucari, C. (2012). Mitochondria as sensors and regulators of calcium signalling. Nature Reviews Molecular Cell Biology 13, 566–578.

Rungta, R. L., Bernier, L.-P., Dissing-Olesen, L., Groten, C. J., LeDue, J. M., Ko, R., … MacVicar, B. A. (2016). Ca^2+^ transients in astrocyte fine processes occur via Ca^2+^ influx in the adult mouse hippocampus. Glia 64, 2093–2103.

Saito, K., Shigetomi, E., Yasuda, R., Sato, R., Nakano, M., Tashiro, K., … Koizumi, S. (2018). Aberrant astrocyte Ca^2+^ signals “AxCa signals” exacerbate pathological alterations in an Alexander disease model. Glia 66, 1053–1067.

Savtchouk, I., & Volterra, A. (2018). Gliotransmission: beyond black-and-white. The Journal of neuroscience: the official journal of the Society for Neuroscience 38, 14–25.

Sharp, A. H., Nucifora, F. C., Blondel, O., Sheppard, C. A., Zhang, C., Snyder, S. H., … Ross, C. A. (1999). Differential cellular expression of isoforms of inositol 1,4,5-triphosphate receptors in neurons and glia in brain. The Journal of comparative neurology 406, 207–20.

Sherwood, M. W., Arizono, M., Hisatsune, C., Bannai, H., Ebisui, E., Sherwood, J. L., … Mikoshiba, K. (2017). Astrocytic IP_3_Rs: contribution to Ca^2+^ signalling and hippocampal LTP. Glia 65, 502–513.

Shigetomi, E., Bushong, E. A., Haustein, M. D., Tong, X., Jackson-Weaver, O., Kracun, S., … Khakh, B. S. (2013). Imaging calcium microdomains within entire astrocyte territories and endfeet with GCaMPs expressed using adeno-associated viruses. The Journal of General Physiology 141, 633–647.

Shigetomi, E., Tong, X., Kwan, K. Y., Corey, D. P., & Khakh, B. S. (2012). TRPA1 channels regulate astrocyte resting calcium and inhibitory synapse efficacy through GAT-3. Nature Neuroscience 15, 70–80.

Srinivasan, R., Huang, B. S., Venugopal, S., Johnston, A. D., Chai, H., Zeng, H., … Khakh, B. S. (2015). Ca^2+^ signaling in astrocytes from Ip3r2^−/−^ mice in brain slices and during startle responses in vivo. Nature Neuroscience 18, 708–717.

Suzuki, J., Kanemaru, K., & Iino, M. (2016). Genetically encoded fluorescent indicators for organellar calcium imaging. Biophysical Journal 111, 1119–1131.

Suzuki, J., Kanemaru, K., Ishii, K., Ohkura, M., Okubo, Y., & Iino, M. (2014). Imaging intraorganellar Ca^2+^ at subcellular resolution using CEPIA. Nature communications 5, 4153.

Szabadkai, G., Bianchi, K., Várnai, P., De Stefani, D., Wieckowski, M. R., Cavagna, D., … Rizzuto, R. (2006). Chaperone-mediated coupling of endoplasmic reticulum and mitochondrial Ca^2+^ channels. The Journal of cell biology 175, 901–11.

Takata, N., Mishima, T., Hisatsune, C., Nagai, T., Ebisui, E., Mikoshiba, K., & Hirase, H. (2011). Astrocyte calcium signaling transforms cholinergic modulation to cortical plasticity in vivo. Journal of Neuroscience 31, 18155–18165.

Takata, N., Nagai, T., Ozawa, K., Oe, Y., Mikoshiba, K., & Hirase, H. (2013). Cerebral blood flow modulation by basal forebrain or whisker stimulation can occur independently of large cytosolic Ca^2+^ signaling in astrocytes. PLoS ONE 8, e66525.

Tamamushi, S., Nakamura, T., Inoue, T., Ebisui, E., Sugiura, K., Bannai, H., & Mikoshiba, K. (2012). Type 2 inositol 1,4,5-trisphosphate receptor is predominantly involved in agonist-induced Ca^2+^ signaling in Bergmann glia. Neuroscience Research 74, 32–41.

Verkhratsky, A., Orkand, R. K., & Kettenmann, H. (1998). Glial calcium: homeostasis and signaling function. Physiological Reviews 78, 99–141.

Volterra, A., & Meldolesi, J. (2005). Astrocytes, from brain glue to communication elements: the revolution continues. Nature Reviews Neuroscience 6, 626–640.

Wang, F., Smith, N. A., Xu, Q., Fujita, T., Baba, A., Matsuda, T., … Nedergaard, M. (2012)(a). Astrocytes modulate neural network activity by Ca^2+^-dependent uptake of extracellular K^+^. Science Signaling 5, ra26.

Wang, F., Xu, Q., Wang, W., Takano, T., & Nedergaard, M. (2012)(b). Bergmann glia modulate cerebellar Purkinje cell bistability via Ca^2+^-dependent K^+^ uptake. Proceedings of the National Academy of Sciences of the United States of America 109, 7911–7916.

Yamamoto-Hino, M., Miyawaki, A., Kawano, H., Sugiyama, T., Furuichi, T., Hasegawa, M., & Mikoshiba, K. (1995). Immunohistochemical study of inositol 1,4,5-trisphosphate receptor type 3 in rat central nervous system. Neuroreport 6, 273–6.

Yang, J., Yang, H., Liu, Y., Li, X., Qin, L., Lou, H., … Wang, H. (2016). Astrocytes contribute to synapse elimination via type 2 inositol 1,4,5-trisphosphate receptor-dependent release of ATP. eLife 5, e15043.

Zhang, Y., Chen, K., Sloan, S. A., Bennett, M. L., Scholze, A. R., O’Keeffe, S., … Wu, J. Q. (2014). An RNA-sequencing transcriptome and splicing database of glia, neurons, and vascular cells of the cerebral cortex. Journal of Neuroscience 34, 11929–11947.

